# Detection and Interspecies Comparison of SARS-CoV-2 Delta Variant (AY.3) in Feces from a Domestic Cat and Human Samples

**DOI:** 10.1101/2022.01.31.478506

**Authors:** Olivia C. Lenz, Andrew D. Marques, Brendan J. Kelly, Kyle G. Rodino, Stephen D. Cole, Ranawaka A.P.M. Perera, Susan R. Weiss, Frederic D. Bushman, Elizabeth M. Lennon

**Author notes:** These first authors contributed equally to this article (Olivia C. Lenz, Andrew D. Marques).

## Abstract

Severe acute respiratory syndrome coronavirus 2 (SARS-CoV-2) infections have spilled over from humans to companion and wild animals since the inception of the global COVID-19 pandemic. However, whole genome sequencing data of the viral genomes that infect non-human animal species has been scant. Here, we detected and sequenced a SARS-CoV-2 delta variant (AY.3) in fecal samples from an 11-year-old domestic house cat previously exposed to an owner who tested positive for SARS-CoV-2. Molecular testing of two fecal samples collected 7 days apart yielded relatively high levels of viral RNA. Sequencing of the feline-derived viral genomes showed the two to be identical, and differing by between 4 and 14 single nucleotide polymorphisms in pairwise comparisons to human-derived lineage AY.3 sequences collected in the same geographic area and time period. However, several mutations unique to the feline samples reveal their divergence from this cohort on phylogenetic analysis. These results demonstrate continued spillover infections of emerging SARS-CoV-2 variants that threaten human and animal health, as well as highlight the importance of collecting fecal samples when testing for SARS-CoV-2 in animals. To the authors’ knowledge, this is the first published case of a SARS-CoV-2 delta variant in a domestic cat in the United States.

## 1. Introduction

Severe acute respiratory syndrome coronavirus 2 (SARS-CoV-2) infections have spilled over from humans to numerous animal species, including domestic cats and dogs, non-domestic large felids, minks, and white-tailed deer, amongst others [1–4]. Several species, including domestic cats, transmit their infection to naive conspecifics under experimental conditions [5]. A number of recent studies have demonstrated natural spillover infections in white-tailed deer (*Odocoileus virginianus*), with likely spread amongst wild deer in the field [2, 6, 7]. Furthermore, human-to-mink and mink-to-human transmission has been documented in mink farms in the Netherlands [4]. These findings provide evidence that SARS-CoV-2 may establish itself in one or more enzootic reservoirs that threaten both non-human animal species and humans.

Emerging SARS-CoV-2 variants have distinct host species ranges. For example, the beta (B.1.351) variant infects deer mice (*Peromyscus* spp.) and laboratory mouse strains, whereas the original strain cannot [8]. Tracking the natural host range of each variant can further clarify potential enzootic reservoir formation and consequential secondary spillover events. This is especially important with the still-widespread delta variant, which transmits readily in humans, can cause severe disease, and is associated with higher rates of vaccine breakthrough infections than lineages that emerged earlier in the epidemic [9–11]. The delta variant encompasses lineages such as B.1.617.2 and AY.3 that share common defining mutations like spike L452R, P681R, and D950N [12, 13].

Thus far one dog in the United States contracted the SARS-CoV-2 delta variant lineage AY.3 [14], and several Asiatic lions in India [15, 16] as well as three domestic cats in China [17] have recently tested positive for the delta variant lineage B.1.617.2.

Here, we report a SARS-CoV-2 delta variant (AY.3) detected in fecal specimens from a domestic house cat in the Delaware Valley region of southeastern Pennsylvania. The animal had a known human COVID-19 exposure and presented to the veterinary hospital for gastrointestinal signs. Whole genome sequencing and phylogenetic analysis revealed lineage AY.3 with several mutations unique among human-derived viral genomes of the same geographic area. To our knowledge, this is the first published case of a SARS-CoV-2 delta variant in a domestic cat in the United States, and the first ever published case of lineage AY.3 in a domestic cat. Our current findings add to the growing body of evidence that further spillover transmission of the delta variant to non-human animals is on-going.

## 2. Materials and Methods

### 2.1 Animal and Human Subjects

Local human-derived viral sequences were gathered as described in a previous publication where sequence data can be accessed [18]. The University of Pennsylvania Institutional Review Board (IRB) reviewed the human research protocol and deemed the limited data elements extracted with positive human SARS-CoV-2 specimens to be exempt from human subject research per 45 CFR 46.104, category 4 (IRB #848605). Informed owner consent was provided for all procedures involving the cat. The University of Pennsylvania Institutional Animal Care and Use Committee (IACUC) and Privately Owned Animal Protocol (POAP) Committee approved the protocol (IACUC/POAP #806977). Consent was obtained from the state animal health officials to collect specimens from the cat for SARS-CoV-2 testing, and for submission of “non-negative” specimens to the National Veterinary Services Laboratory (Ames, IA) for confirmation of a positive test.

### 2.2 SARS-CoV-2 Clinical Testing

RNA was extracted from specimens using a QIAamp Viral RNA Mini Kit (Qiagen, Germantown, MD). Testing for SARS-CoV-2 was performed at the university microbiology laboratory using the CDC 2019 Novel Coronavirus (2019-nCoV) Real-Time Reverse Transcriptase (RT)–PCR Diagnostic Panel (IDT, Coralville, IA). The university microbiology laboratory is a member laboratory of the Food and Drug Administration (FDA) Veterinary Laboratory Investigation and Response Network (Vet-LIRN). As part of this network, the university microbiology laboratory completed an Inter-Laboratory Comparison Exercise (ICE) of SARS-CoV-2 Molecular Detection Assays Being Used by Veterinary Diagnostic Laboratories in August 2020.

### 2.3 SARS-CoV-2 Whole Genome Sequencing

The POLAR protocol was used for sequencing genomes [19]. Specifically, 5 μl of viral RNA, 0.5μl of 10mM dNTPs Mix (Thermo Fisher, 18427013), 0.5 μl of 50 μM Random Hexamers (Thermo Fisher, N8080127), and 1 μl water was heated at 65°C for 5 minutes. Reverse transcription was performed with a reaction containing 6.5 μl from the previous step, 0.5 μl of RNaseOut (Thermo Fisher, 18080051), 0.5 μl of 0.1M DTT (Thermo Fisher, 18080085), 0.5 μl SuperScript III Reverse Transcriptase (Thermo Fisher, 18080085), and 2 μl of 5X First-Strand Buffer (Thermo Fisher, 18080085). This mixture was heated at 42°C for 50 minutes, then incubated at 70°C for 10 minutes. ARTIC-nCoV2019 version 4 primers were used (IDT) to amplify the product by PCR in a reaction containing 2.5 μl of the product from the previous step, 0.5 μl of 10 mM dNTPs Mix (NEB, N0447S), either 4.0 μl of primer set 1 or 3.98 μl of primer set 2, 0.25 μl Q5 Hot Start DNA Polymerase (NEB, M0493S), 5 μl of 5X Q5 Reaction Buffer (NEB, M0493S), and water to bring to 25 μl. The mixture was amplified with 1 cycle at 98°C for 30 seconds, then 25 cycles at 98°C for 15 seconds and 65°C for 5 minutes. Products from primer set 1 and 2 were combined and then brought to a concentration of 0.25 ng/μl. The Nextera XT Library Preparation Kit (Illumina, FC-131-1096) and the IDT for Illumina DNA/RNA UD Indexes (Illumina, 20027213, 20027214, 20027215, 20027216) were used for library prep. Each sample was quantified with the Quant-iT PicoGreen dsDNA quantitation kit (Invitrogen, P7589). The samples were then pooled and sequenced on an Illumina NextSeq.

### 2.4 SARS-CoV-2 Whole Genome Sequencing

Sequences were trimmed and aligned to the Wuhan reference sequence (NC_045512.2). Alignment used the BWA aligner tool (v0.7.17) [20]. Samtools package (v1.10) was used to remove reads that did not align to the reference [21]. To accept a genome as high quality, we required that coverage must be ≥ 5 read depth for ≥ 95% of the genome. The Bcftools package (v1.10.2-34) was used to call the variant positions [22]. The Pangolin lineage software (Pangolin version 3.1.17 with the PangoLEARN 2021-12-06 release) was used to assign variants. A pipeline developed by Everett et al. was used to assign point mutations [23].

### 2.5 SARS-CoV-2 Whole Genome Sequencing

To construct phylogenetic trees, NextClade was used for alignment [24], IQ-Tree (v1.6.12) was used to generate the phylogenetic tree [25–28], and FigTree v1.4.4 was used to visualize the tree.

## 3. Results

### 3.1. Case Description

In September 2021, an 11-year-old indoor-only female spayed domestic shorthair cat (*Felis catus)* was presented to the Ryan Veterinary Hospital Emergency Service at the University of Pennsylvania School of Veterinary Medicine following several days of anorexia, lethargy, soft stools, and vomiting as well as a known COVID-19 exposure. One of the cat’s owners tested positive for SARS-CoV-2 prior to onset of the cat’s clinical signs. At the time of sample collection, the cat had been isolated from the infected human for 11 days and was cared for by another household member who repeatedly tested negative.

The cat had a medical history of presumptive chronic enteropathy, which had been successfully managed with a hydrolyzed protein diet and for which further diagnostics were not performed, as well as hypertrophic obstructive cardiomyopathy that was treated with atenolol.

On physical examination, the cat’s heart rate, respiratory rate, and temperature were within normal limits, with normal lung sounds on cardiothoracic auscultation. She was mildly uncomfortable on abdominal palpation. The remainder of her physical examination was un-remarkable.

A fecal sample was submitted for polymerase chain reaction (PCR) testing for infectious agents associated with feline gastrointestinal disease: Feline *parvovirus, Tritrichomonas foetus, Campylobacter jejuni/coli, Cryptosporidium* spp., *Cryptosporidium felis, Salmonella* spp., *Giardia* spp., *Clostridium difficile* toxin A/B, and *Clostridium perfringens* enterotoxin. All tests were negative.

### 3.2 Molecular Detection and Sequencing

The fecal sample was tested for SARS-CoV-2 using the Centers for Disease Control 2019 Novel Coronavirus real time PCR (RT-PCR) Diagnostic Panel. The sample tested positive for both viral nucleocapsid targets with cycle threshold (Ct) values of 26.3 and 27.7. The oropharyngeal swab sample was negative. To comply with reportable disease mandates, an aliquot of the fecal sample was sent to the National Veterinary Services Laboratory (NVSL) (Ames, IA) and confirmed as positive. A second fecal sample collected seven days later was positive with Ct values of 27.7 and 28. Attempts to isolate replication-competent virus were unsuccessful.

We performed SARS-CoV-2 whole genome sequencing (WGS) on the two samples from the cat. We received 99.7% and 98.3% coverage with a mean coverage of 1,843 and 374 reads for the two samples respectively [18]. WGS performed by NVSL yielded nearly identical results using slightly different techniques. Differences between our groups’ sequencing results are attributed to differences in primers used at the time of sequencing [29].

### 3.3 Comparison to Known Sequences in the Delaware Valley

The feline-derived SARS-CoV-2 genome was identified as delta variant lineage AY.3. The sequences obtained from the fecal specimens on days 1 and 8 were identical, and therefore stable over a 7-day period. In addition to the mutations associated with known human-derived AY.3 sequences, our sample has several that are uncommon or unique (Table 1). Out of over 4,200 human samples that we have sequenced from our geographic region, the Delaware Valley in Pennsylvania, 10 single nucleotide polymorphisms (SNPs) found in the feline-derived samples have been identified in less than 5% of them (“Percent in Human Dataset” column of Table 1). 7 of these 10 nucleotide mutations were silent mutations. The 3 rarer non-silent mutations include an I3731V mutation in ORF1ab (Nsp6 protein), N2426T mutation in ORF1ab (Nsp16 protein), and D80N in Spike.

**Table 1.** Mutation table outlining the 45 mutations detected in the feline fecal samples. Both samples collected from the cat (VSP3509 and VSP3510) contained the same mutations. These mutations are compared to the random dataset consisting of 4,250 human-derived genomes representing the geographical area of residence for the cat described here. The original Wuhan isolate (NC_045512.2) was used as a reference.

| Genomic Position | Gene | Affected Protein | Protein Mutation | Nucleotide Mutation | Reference Nucleotide | Percent in Human Dataset |
| --- | --- | --- | --- | --- | --- | --- |
| 210 | intergenic |  |  | T | G | 40% |
| 241 | intergenic |  |  | T | C | 99.6% |
| 3037 | ORF1ab | nsp3 | silent | T | C | 99.95% |
| 4181 | ORF1ab | nsp3 | A1306S | T | G | 34.64% |
| 6402 | ORF1ab | nsp3 | P2046L | T | C | 34.73% |
| 7124 | ORF1ab | nsp3 | P2287S | T | C | 34.64% |
| 8140 | ORF1ab | nsp3 | silent | T | C | 1.46% |
| 8986 | ORF1ab | nsp4 | silent | T | C | 34.68% |
| 9053 | ORF1ab | nsp4 | V2930L | T | G | 34.66% |
| 9080 | ORF1ab | nsp4 | silent | T | C | 0.02% |
| 10029 | ORF1ab | nsp4 | T3255I | T | C | 38.16% |
| 11201 | ORF1ab | nsp6 | T3646A | G | A | 34.68% |
| 11332 | ORF1ab | nsp6 | silent | G | A | 34.71% |
| 11456 | ORF1ab | nsp6 | I3731V | G | A | 4.66% |
| 14408 | ORF1ab | nsp12 (RdRp) | P314L | T | C | 99.53% |
| 14520 | ORF1ab | nsp12 (RdRp) | silent | T | C | 0% |
| 15451 | ORF1ab | nsp12 (RdRp) | G662S | A | G | 39.36% |
| 16466 | ORF1ab | nsp13 (Hel) | P1000L | T | C | 39.32% |
| 19220 | ORF1ab | nsp14 (ExoN) | A1918V | T | C | 34.59% |
| 20744 | ORF1ab | nsp16 (2'-O-MT) | N2426T | C | A | 0% |
| 21618 | S | spike | T19R | G | C | 39.86% |
| 21800 | S | spike | D80N | A | G | 0% |
| 21987 | S | spike | G142D | A | G | 13.04% |
| 22029 | S | spike | del 6 | delAGTTCA | GAGTTCA | 39.13% |
| 22917 | S | spike | L452R | G | T | 41.58% |
| 22995 | S | spike | T478K | A | C | 40.45% |
| 23284 | S | spike | silent | C | T | 2.47% |
| 23403 | S | spike | D614G | G | A | 99.98% |
| 23604 | S | spike | P681R | G | C | 40.07% |
| 24410 | S | spike | D950N | A | G | 40.02% |
| 25339 | S | spike | silent | T | C | 2.64% |
| 25469 | ORF3a | ORF3a | S26L | T | C | 40.05% |
| 26767 | M | membrane | I82T | C | T | 41.55% |
| 27638 | ORF7a | ORF7a | V82A | C | T | 39.29% |
| 27752 | ORF7a | ORF7a | T120I | T | C | 39.6% |
| 27874 | ORF7b | ORF7b | T40I | T | C | 34.42% |
| 28248 | ORF8 | ORF8 | del 6 | delGATTTC | AGATTTC | 38.87% |
| 28271 | intergenic |  | del 1 | delA | TAAAA | 62.19% |
| 28461 | N | nucleocapsid | D63G | G | A | 39.41% |
| 28881 | N | nucleocapsid | R203M | T | G | 39.91% |
| 28916 | N | nucleocapsid | G215C | T | G | 34.49% |
| 29050 | N | nucleocapsid | silent | A | G | 4.64% |
| 29402 | N | nucleocapsid | D377Y | T | G | 42.21% |
| 29509 | N | nucleocapsid | silent | T | C | 4.78% |
| 29742 | intergenic |  |  | T | G | 37.01% |

The feline-derived sample (VSP3509) differed by between 4 and 14 SNPs in pairwise comparisons with human samples drawn from a random sampling of human-derived lineage AY.3 sequences from the Delaware Valley collected between 6/21/2021 and 11/18/2021 (Figure 1).

**Figure 1.**
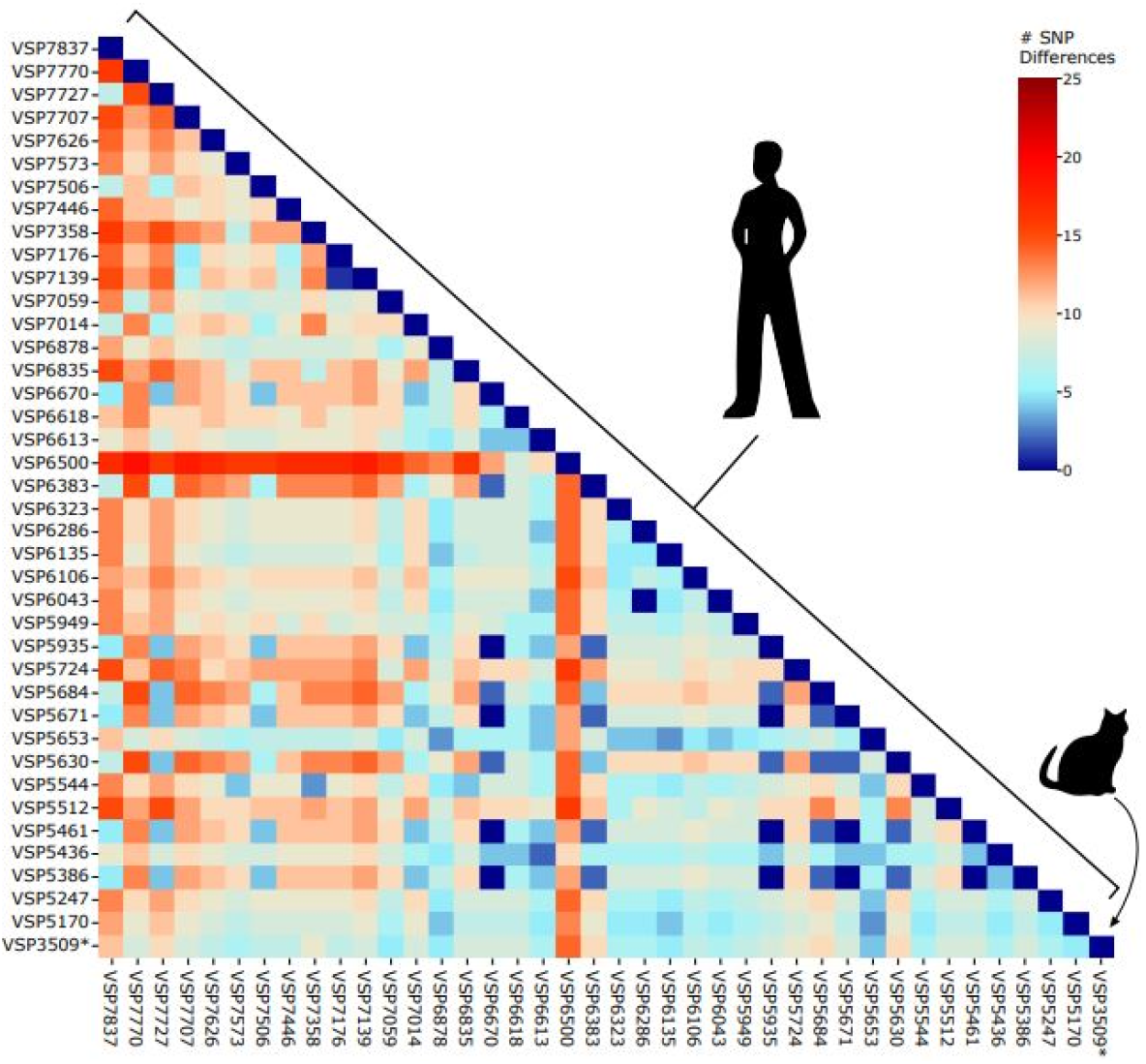
Pairwise distances between AY.3 sequences in the Delaware Valley. Included are the feline-derived sequence (VSP3509) and human-derived sequences. The number of SNPs separating each pair of lineages is shown by the color code (key to the right of the figure).

Phylogenetic analysis reveals that the cat-derived sequence, as well as another feline-derived SARS-CoV-2 lineage AY.3 genome found on GISAID, is divergent from the human sequences (Figure 2). Therefore, while there are few SNPs that differentiate the cat-derived samples from the human-derived samples nearest in sequence, the unique SNPs (Table 1) cause the cat samples to appear more distant on the phylogenetic tree. Some of these mutations may be enriched in samples from cats, however a larger dataset is necessary to draw this conclusion.

**Figure 2.**
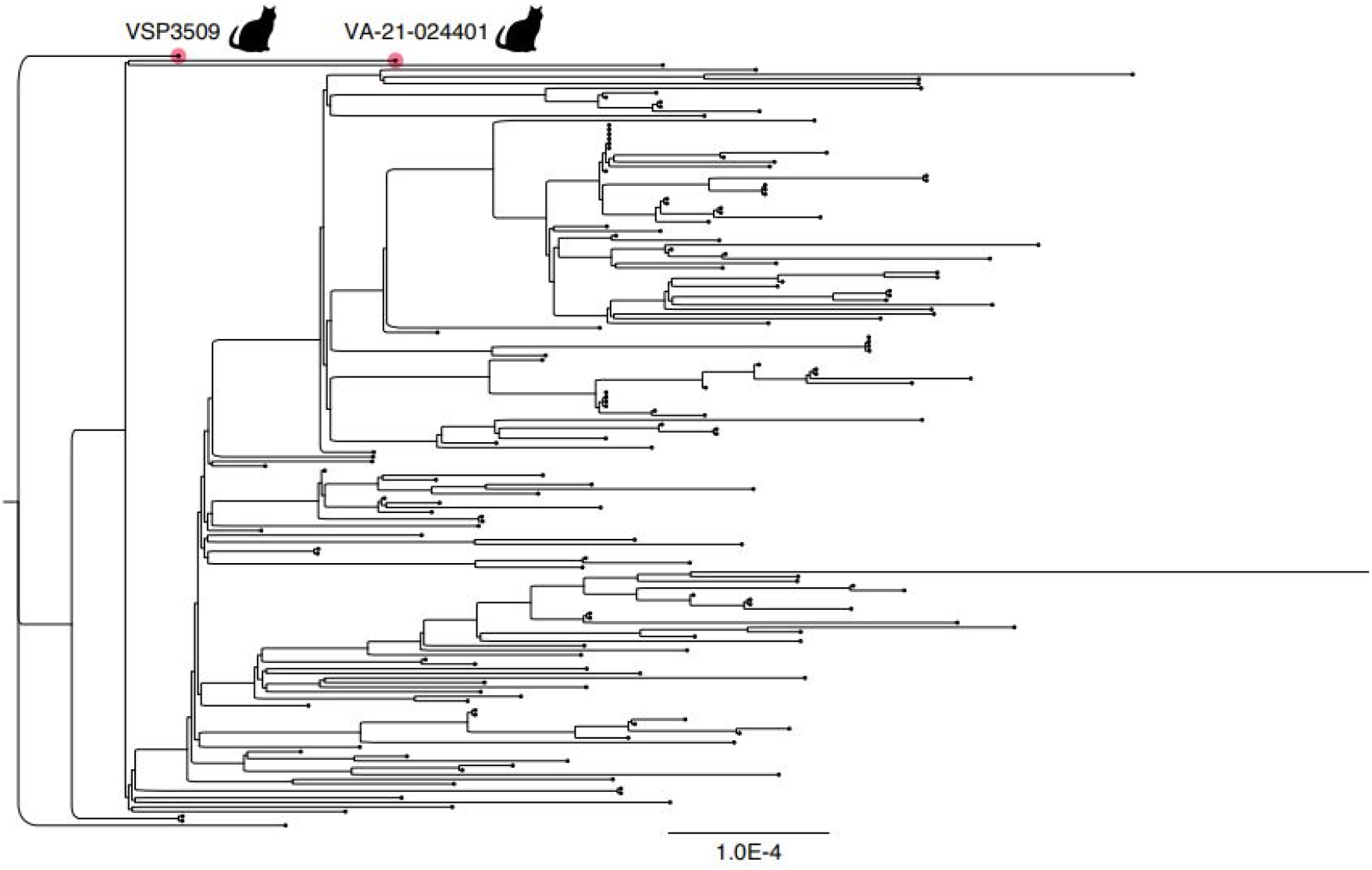
Phylogenetic tree depicting the distances of AY.3 genomes. Included are the cat specimen discussed in this article in addition to an AY.3 cat-derived genome previously collected on 8/5/2021 in Virginia, USA (EPI_ISL_5761527) compared to a random sampling of SARS-CoV-2 human-derived genomes in the Delaware Valley.

## 4. Discussion

To date, published reports on the SARS-CoV-2 delta variant lineage AY.3 have described infection of humans and one domestic dog [14–16, 18]. Here, we report delta variant lineage AY.3 in fecal samples from a domestic cat who was exposed to a human with SARS-CoV-2.

Two feline fecal samples collected seven days apart both had Ct values between 26-28, quantities sufficient for WGS, indicating relatively high levels of genomic replication. Furthermore, the cat had been isolated from the infected owner for 11 days and 18 days by the dates of the first and second positive SARS-CoV-2 tests, respectively, reducing likelihood that the cat sample was falsely positive (for example, due to pass-through contamination from the infected owner during self-grooming). However, we cannot determine whether the cat’s clinical signs are attributable to COVID-19, a flare-up of chronic enteropathy, or a combination. Anorexia, diarrhea, and vomiting are among the clinical signs observed in feline patients who test positive for SARS-CoV-2 by RT-PCR on fecal samples [30]. Prior to the COVID-19 exposure, however, the cat’s enteropathy had been managed successfully with a prescription diet for months with no clinical signs.

The discovery of a delta variant lineage AY.3 sequence in a feline sample, taken together with detection of delta variant lineage B.1.617.2 in non-human animal species, suggests that interspecies transmission of SARS-CoV-2 occurs among multiple delta variants. Recently, identical sequences of lineage B.1.575 were discovered in a pet dog and cat and their owner [31], demonstrating that minimal viral evolution is required to overcome species barriers in at least one variant. Because we do not have the infected owner’s SARS-CoV-2 sequence, we cannot determine whether the mutations found in the feline-derived sequence originate from the presumptive infective human, or whether they arose with the species barrier jump.

While fecal samples from the infected cat contained relatively high levels of viral genetic material, SARS-CoV-2 was not detected on the oropharyngeal swab collected on the day of presentation to the veterinary hospital. This has been reported once previously in companion animals [31]. Transmission and pathophysiology appear to differ among species and may be responsible for this discrepancy, although one study found that over half of human patients infected with SARS-CoV-2 continued to test positive on fecal samples for approximately 11 days after respiratory tract samples tested negative [32]. Therefore, we may have missed the window for detecting SARS-CoV-2 in respiratory samples from our feline patient. Regardless, our data underscores the importance of taking fecal samples in addition to oropharyngeal or nasal swabs for maximal sensitivity when testing for the virus in non-human animals.

Since domestic felines can support relatively efficient replication of SARS-CoV-2 viral genomes similar to those that infect humans, can transmit SARS-CoV-2 viruses to naïve conspecifics, and frequently have a high degree of contact with humans, they have the potential to become an enzootic reservoir for the virus. Cat population dynamics contribute to this potential, as owned indoor-outdoor cats may mingle with each other as well as free-roaming unowned cats and various wildlife species, creating an unseen network between households, free-roaming community cats, and wildlife populations. Transmission and reservoir formation of SARS-CoV-2 in any non-human animal species poses a threat to domestic animal, wildlife, and human health. This highlights the need to closely track SARS-CoV-2 variants of concern in domestic house cats to better understand the intertwined nature of animal and human health in this global pandemic.

## Author Contributions

Conceptualization, E.M.L, F.D.B., O.C.L., A.D.M.; methodology, A.D.M., B.J.K, K.G.R., S.D.C., R.A.P.M.P., S.R.W.; formal analysis, A.D.M., F.D.B.; data curation, A.D.M.; writing—original draft preparation, O.C.L., A.D.M., E.M.L., F.D.B.; writing—review and editing, O.C.L., A.D.M., B.J.K., K.G.R., S.D.C., R.A.P.M.P., S.R.W., F.D.B., E.M.L. All authors have read and agreed to the published version of the manuscript.

## Institutional Review Board Statement

Ethical review and approval were waived for this study. The University of Pennsylvania Institutional Review Board (IRB) reviewed the human research protocol and deemed the limited data elements extracted with positive human SARS-CoV-2 specimens to be exempt from human subject research per 45 CFR 46.104, category 4 (IRB #848605). Informed owner consent was provided for all procedures in-volving the cat. The University of Pennsylvania Institutional Animal Care and Use Committee (IACUC) and Privately Owned Animal Protocol (POAP) Committee approved the protocol (IACUC/POAP #806977).

## Informed Consent Statement

Not applicable.

## Data Availability Statement

The cat-derived viral genome sequences acquired in this study have been deposited in GISAID under accession numbers EPI_ISL_8599342 and EPI_ISL_8599343.

## Acknowledgments

We are grateful to individuals and their families who volunteered to provide specimens, to Laurie Zimmerman for artwork, to Jaclyn Dietrich and Elias Braun for technical support, and to Dr. Leslie King for manuscript editing. We acknowledge help from all the staff of the Philadelphia Department of Public Health.

